# Monosaccharides Drive *Salmonella* Gut Colonization in a Context-Dependent Manner

**DOI:** 10.1101/2024.08.06.606610

**Authors:** Christopher Schubert, Bidong D. Nguyen, Andreas Sichert, Nicolas Näpflin, Anna Sintsova, Lilith Feer, Jana Näf, Benjamin B.J. Daniel, Yves Steiger, Christian von Mering, Uwe Sauer, Wolf-Dietrich Hardt

**Affiliations:** Institute of Microbiology, Department of Biology, ETH Zurich, Zurich, Switzerland; Institute of Molecular Systems Biology, ETH Zurich, Zurich, Switzerland; Department of Molecular Life Sciences and Swiss Institute of Bioinformatics, University of Zurich, Switzerland

## Abstract

The carbohydrates that fuel gut colonization by *S*. Typhimurium are not fully known. To investigate this, we designed a quality-controlled mutant pool to probe the metabolic capabilities of this enteric pathogen. Using WISH-barcoding, we tested 35 metabolic mutants across five different mouse models, allowing us to differentiate between context-dependent and context-independent nutrient sources. Results showed that *S*. Typhimurium uses D-glucose, D-mannose, D-fructose, and D-galactose as context-independent carbohydrates across all models. The utilization of N-acetylglucosamine and hexuronates, on the other hand, was context-dependent. Furthermore, we showed that D-fructose is important in strain-to-strain competition between *Salmonella* serovars. Complementary experiments confirmed that D-glucose, D-fructose, and D-galactose are excellent niches for *S*. Typhimurium to exploit during colonization. Quantitative measurements revealed sufficient amounts of D-glucose and D-galactose in the murine cecum to drive *S*. Typhimurium colonization. Understanding these key substrates and their context-dependent use by enteric pathogens will inform the future design of probiotics and therapeutics to prevent diarrheal infections such as non-typhoidal salmonellosis.

## Introduction

The gut microbiota provides colonization resistance against invading pathogens through diverse mechanisms, including nutrient competition, known as exploitation, and the release of antimicrobial metabolites and type VI secretion systems, termed interference (Caballero-Flores et al., 2023). Despite these mechanisms, *Salmonella enterica* serovar Typhimurium (*S.* Typhimurium), a Gram-negative enteric pathogen, can overcome these defenses, leading to foodborne diarrhea in humans. *S. Typhimurium* is responsible for a considerable number of diarrheal infections (EFSA, 2022). Through animal infection experiments, it has been observed that gut colonization by *S.* Typhimurium occurs in distinct phases (Wotzka et al., 2017). Following oral exposure, *S.* Typhimurium undergoes initial growth without disturbing the resident microbiota (Nguyen et al., 2020). Subsequently, invasion of gut tissue triggers immune responses that decrease the pathogen’s tissue burden and alter the gut-luminal environment. *S.* Typhimurium can then flourish in the inflamed gut lumen by utilizing anaerobic and microaerophilic respiration (Rivera-Chavez et al., 2016; Winter et al., 2010; Winter et al., 2013). In recent years, some metabolic strategies have been clarified, particularly regarding the types of electron acceptors and some of the electron donors available for *S*. Typhimurium to be utilized (Faber et al., 2017; Gillis et al., 2018; Gillis et al., 2019; Humphries et al., 2003; Jones et al., 2007; Jones et al., 2011; Litvak et al., 2019; Lopez et al., 2015; Lopez et al., 2012; Maier, Barthel, et al., 2014; Maier et al., 2013; Nguyen et al., 2020; Rivera-Chavez et al., 2016; Spiga et al., 2017; Stecher et al., 2007; Thiennimitr et al., 2011; Winter et al., 2023; Winter et al., 2010; Winter et al., 2013). Much less is known about carbohydrate utilization. Earlier work found that *S*. Typhimurium uses L-arabinose and the Amadori product fructose-asparagine to proliferate in the inflamed gut (Ali et al., 2014). Additionally, it has been demonstrated that the hexitol D-galactitol plays a context-dependent role in inter- and intraspecies competition (Eberl et al., 2021; Gül et al., 2023). However, a key limitation of this previous research is the predominant use of a single mouse model, which may introduce bias and limit the observable context-dependency of carbohydrate use in these colonization experiments.

Transposon mutagenesis provides a powerful technology to identify mutants with fitness defects during animal infections (Bourgeois & Camilli, 2023; Chaudhuri et al., 2013; Fong et al., 2023; Hensel et al., 1995; Wang et al., 2024). Genome-wide random-barcoded transposon sequencing (RB-TnSeq) has uncovered metabolic strategies used by *S*. Typhimurium to invade and thrive in the gut (Nguyen et al., 2020). This method creates large pools of mutants with unique barcodes for colonization studies but poses bioinformatic challenges, requiring many mice and high sequencing costs. These technical hurdles hinder systematic comparisons between different mouse models. Additionally, the target gene of many mutants is not effectively deactivated, or genes of interest may be missing from the pool, requiring more mutants and mice for statistically significant data. Our study adopts a different approach, leveraging extensive knowledge of *S*. Typhimurium and *E. coli* central carbohydrate metabolism (Karp et al., 2023; Mayer & Boos, 2005). We created a pool of uniquely WISH-barcoded (Daniel et al., 2024), rationally selected *S*. Typhimurium mutants in carbohydrate utilization to represent the metabolic capacity.

Colonization resistance, as highlighted by (Campbell et al., 2023), points out the pivotal role of the microbiome. This protection is achieved by a diverse microbiota depleting available nutrients and is enhanced by the presence of taxa closely related to the incoming pathogen. For instance*, E. coli* increases protection against *Salmonella* infection (Brugiroux et al., 2016; Velazquez et al., 2019; Wotzka et al., 2019). In many cases, the crucial factor for this protection is the metabolic resource overlap between the microbiota, key species, and the invading pathogen (Spragge et al., 2023). Therefore, understanding the metabolic systems used by *S*. Typhimurium in various microbiota contexts is essential. We studied the same *S*. Typhimurium mutant pool across five mouse models: germ-free C57BL/6J, streptomycin-pretreated C57BL/6J (SPF) and 129S6/SvEvTac (SPF), low-complexity microbiome (LCM) C57BL/6J, and oligo mouse microbiota (OligoMM^12^) C57BL/6J mouse models (**Fig. 1A**). Specifically, the LCM and OligoMM^12^ models provide a balanced approach to studying pathogen colonization and interactions with the host microbiota, featuring partial colonization resistance that simulates natural infection kinetics. This study aimed to identify carbohydrate sources essential for *S*. Typhimurium colonization and examine their context dependency.

**Figure 1.**
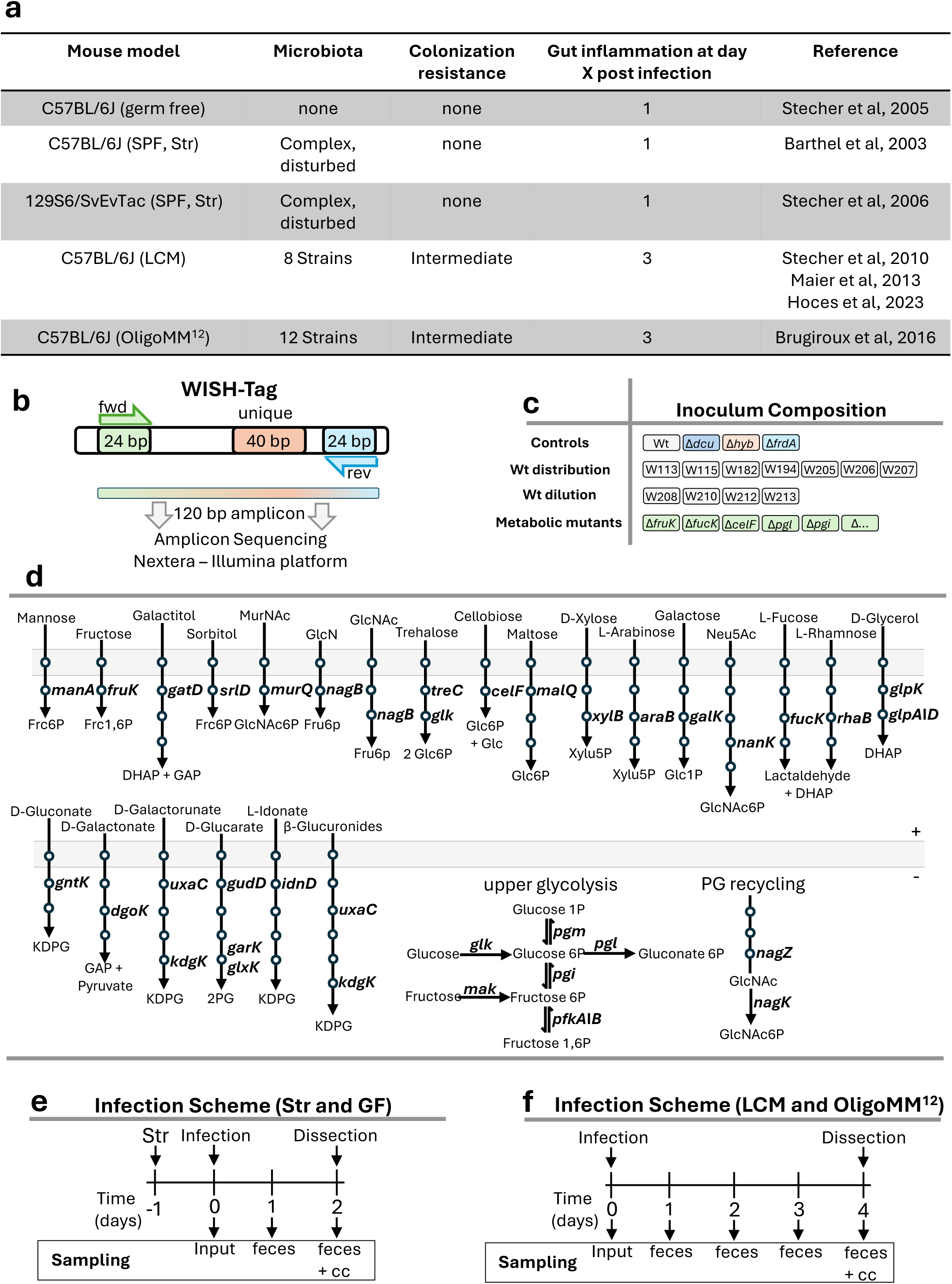
Rational design of a WISH-barcoded *S*. Typhimurium mutant pool. **A** Overview of the five mouse models used in this study, showing microbiota composition, colonization resistance, and when gut inflammation typically occurs. **B** Schematic representation of the wild type isogenic standardized (WISH)-tag (Daniel et al., 2024). **C** Composition of the inoculum pool, as detailed in the main text. **D** A schematic depiction illustrates the cytosolic genes chosen for the mutant pool. Within the respective pathway, only the designated gene is emphasized, while each circle denotes an enzymatic step in the degradation of the corresponding carbohydrate. Refer to **Table S12** for abbreviations. **E** Mouse infection protocol and sampling procedures for the germ free (GF) and streptomycin (Str) pretreated C57BL/6J and 129S6/SvEvTac mouse models. **F** Mouse infection protocol and sampling procedures for the gnotobiotic low complex microbiota (LCM) and oligo mouse microbiota (OligoMM^12^) model.

## Results

### Rationale for designing a *S*. Typhimurium carbohydrate mutant pool

A rationally designed mutant pool was constructed to study *S*. Typhimurium carbohydrate utilization across different mouse models, using *Salmonella enterica* subsp. 1 serovar Typhimurium SL1344 (Hoiseth & Stocker, 1981). We investigated 35 metabolic mutants using the WISH-barcoding approach (Daniel et al., 2024), which allows quantitative PCR and amplicon sequencing for quantification of the strain abundance (**Fig. 1B, Table S1**). The inoculum comprised four groups: control mutants with known colonization defects, seven wild-type strains to assess the evenness of distribution, a wild-type dilution series, and 35 metabolic mutants deficient in carbohydrate-utilizing enzymes (**Fig. 1C**). The mutants, targeting carbohydrate-specific enzymes like kinases, dehydrogenases, or isomerases, provided insight into carbohydrate utilization without transporter specificity issues. This approach allowed us to systematically analyze the fitness of each mutant in various gut environments, enabling the identification of context-independent and -dependent key carbohydrates essential for *S*. Typhimurium colonization (**Fig. 1D**). Four WISH-barcoded strains were included as controls. Firstly, a Δ*dcuA ΔdcuB ΔdcuC* triple mutant (abbreviated: Δ*dcuABC*) and Δ*frdABCD* (abbreviated Δ*frd*), which encode the C4-dicarboxylate transporters and fumarate reductase, respectively, both crucial for fumarate respiration during gut-luminal growth in LCM mice (Nguyen et al., 2020) and during inflammation (Yoo et al., 2024). Additionally, a Δ*hyb* mutant, encoding a hydrogenase important for utilizing hydrogen as an electron donor (Maier et al., 2013) and finally a SL1344 wild type was also included. To assess bottleneck severity, seven SL1344 wild-type strains were used to calculate the Shannon evenness score. A drop below 0.9 indicated a significant bottleneck, potentially leading to false positives; therefore, such samples were excluded (Maier, Diard, et al., 2014). A wild-type dilution series established the measurement window for each mouse and time point, ensuring accurate quantification. Each strain was WISH-barcoded for subsequent abundance quantification in fecal or cecum samples by amplicon sequencing, which was analyzed using the mBARq software (Sintsova et al., 2024).

We employed two primary infection protocols in our experiments (**Fig. 1E-F**). For SPF C57BL/6J and 129S6/SvEvTac mouse models, mice were pre-treated with streptomycin to reduce resident microbiota and achieve consistent SL1344 infection (Barthel et al., 2003; Stecher et al., 2006). We also used gnotobiotic mouse models with a C57BL/6J background: the low-complexity microbiota (LCM) model with 8 bacterial strains (Hoces et al., 2023; Maier et al., 2013; Stecher et al., 2010) and the oligo mouse microbiota model (OligoMM^12^) with 12 strains (Brugiroux et al., 2016). Germ-free mice, lacking any microbiome, were also included (Stecher et al., 2005). These non-SPF models were infected without antibiotic pretreatment. Due to the severity of SL1344 infection and population bottlenecks occurring during the late stage of infection (Maier, Diard, et al., 2014), germ-free and streptomycin-pretreated models were limited to a 2-day infection period. In contrast, infection of gnotobiotic animals could be extended to 4 days, as these models feature much less pronounced gut-luminal inflammation-associated bottlenecks. An inoculum size of 5×10^7^ CFUs ensured each strain was represented by over 100,000 cells, minimizing stochastic effects.

### *S.* Typhimurium pool data was verified across all mouse models

The *S*. Typhimrium infection was monitored by plating feces and cecum content on selective MacConkey plates to quantify bacterial loads. As expected, gut colonization kinetics differed between models. In germ-free and streptomycin-pretreated mice, SL1344 reached 10^10^ CFUs per gram of stool within one day due to reduced microbiota or its absence (**Fig. S1A-C**). In LCM and OligoMM^12^ gnotobiotic models, colonization increased more slowly, saturating at day 3 and 4 post-infection, respectively (**Fig. S1D-E**). The Shannon evenness scores for the seven SL1344 wild type controls were close to 1 for the majority of samples in germ-free mice, streptomycin-pretreated mice, and the LCM mouse model (**Fig. S1F-I**). The OligoMM^12^ model showed more variation, with some samples falling below the 0.9 threshold by day 4 post infection (**Fig. S1J**). Samples with Shannon evenness scores below 0.9 were excluded from further analysis, ensuring high-quality fitness data for all models at all time points.

The WISH-barcoded SL1344 controls deficient in fumarate respiration (Δ*frd* and Δ*dcuABC*) showed 10 to 100-fold attenuation in all models compared to the wild type from day 1 post-infection (**Fig. S1K-O**). Fumarate respiration can function independently of external fumarate by using monosaccharides or L-aspartate, making it the sole option for initial growth when external inorganic electron acceptors are absent (Nguyen et al., 2020; Schubert & Unden, 2022). The Hyb-deficient strain showed no attenuation and sometimes better growth in germ-free and streptomycin-pretreated models but exhibited 5 to 100-fold attenuation in LCM and OligoMM^12^ models (**Fig. S1K-M** compared to **N-O**). The LCM and OligoMM^12^ model highlighted the importance of *hyb* for *S*. Typhimurium colonization, aligning with the expectation that a perturbed microbiota does not provide sufficient hydrogen to drive H_2_/fumarate respiration (Fischbach & Sonnenburg, 2011; Flint et al., 2008; Hoces et al., 2022). These findings validate the experimental data, as previously published for LCM animals (Maier et al., 2013; Nguyen et al., 2020), and newly highlight the context-independent importance of fumarate respiration and the context-dependency of hydrogen utilization in *S*. Typhimurium colonization.

### D-fructose, D-galactose, and D-mannose utilization are crucial for colonization

Our SL1344 pool reveals both context-independent and context-dependent carbohydrate sources that are crucial for *S*. Typhimurium. In **Table S2-S6**, the competitive indices of all 35 metabolic mutants across five mouse models are listed. SL1344 mutants lacking *pgi* and *pgm* consistently fall below detection limits, underscoring the importance of glucose 1-phosphate, glucose 6-phosphate, and fructose 6-phosphate interconversion. The key enzyme of glycolysis, *pfkA*, which encodes 6-phosphofructokinase, was also important across all animal models; however, it was consistently within the range of the detection limit. Additionally, the genes *fruK* (D-fructose), *galK* (D-galactose), and *manA* (D-mannose), which encode 1-phosphofructokinase, galactokinase, and mannose-6-phosphate isomerase respectively, are noteworthy. The *fruK*-deficient SL1344 mutant shows attenuation across all mouse models, with competitive index changes from 10 to 1000-fold (**Fig. 2A-E**). In streptomycin-pretreated models, the competitive index of *fruK* is near 1 on day 1, dropping 10-fold by day 2 post infection (**Fig. 2B-C**), unlike other models where attenuation is seen on day 1 (**Fig. 2A, D-E**). Similarly, the *manA* mutant is attenuated in all models, with a significant drop below the limit of detection in almost all animals in the streptomycin-pretreated C57BL/6J model by day 1 post-infection (**Fig. 2B**). The *galK*-deficient mutant shows context-dependent attenuation only in germ-free and streptomycin-pretreated models (**Fig. 2A-C**). These findings suggest a context-independent role of D-fructose and D-mannose utilization pathways in supporting *S*. Typhimurium growth in the murine gut, while the importance of D-galactose appears to be context-dependent in microbiota-perturbed mouse models.

**Figure 2.**
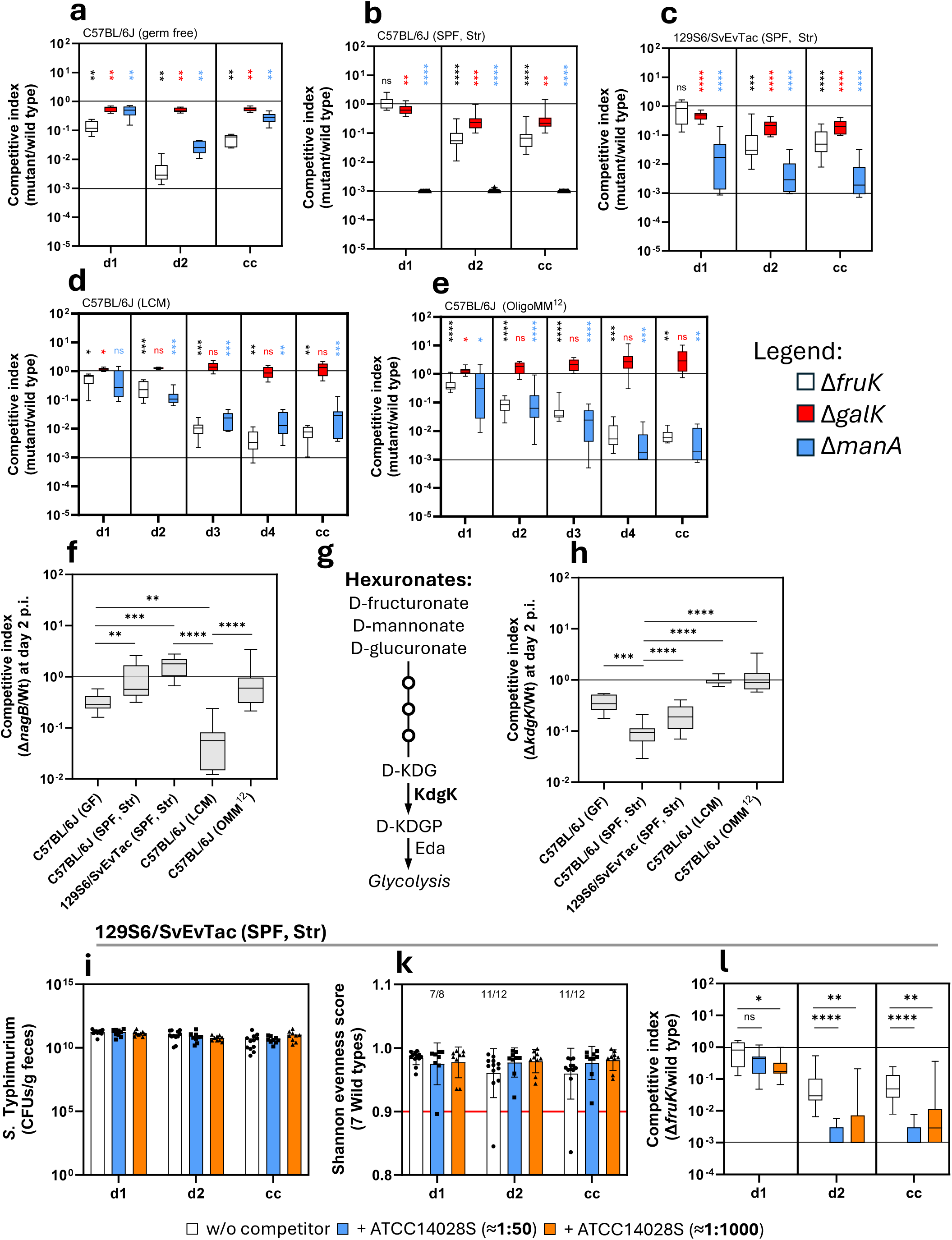
*S*. Typhimurium mutants deficient in D-fructose, D-mannose, and D-galactose are attenuated. **A-E** Competitive index (CI) for D-fructose (Δ*fruK*; white), D-galactose (Δ*galK*; red), and D-mannose (Δ*manA*; blue) in the five different mouse models, as indicated above the diagram. The upper black line indicates wild-type CI of 1 and the lower black line the limit of detection at 10^-3^. The metabolic mutants were statistically compared to the SB300 wild type in the control group. **F** Competitive indices for N-acetylglucosamine (Δ*nagB*) for all five mouse models are indicated on the x-axis. **G** Schematic representation of hexuronate degradation, wherein KdgK catalyses the penultimate step preceding the Entner-Doudoroff pathway (Eda). **H** Competitive indices for hexuronate (Δ*kdgK*) in all five mouse models are indicated on the x-axis. **I** The WISH-barcoded *S*. Typhimurium mutant pool was competed with *S.* Typhimurium ATCC14028S in a 1:50 (mice n=9) and 1:1000 ratio (mice n=9, at least two independent experiments). The SL1344 loads are plotted in CFUs per gram feces in absence and presence of ATCC14028S at both ratios. **K** Shannon evenness score (SES) was calculated for the 7 WISH-barcoded SL1344 wild types in absence and presence of ATCC14028S at both ratios. **L** The competitive index of the SL1344 *fruK* mutant is shown, in absence and presence of the competitor ATCC14028S at both ratios. All competitive experiments are presented in a box-and-whiskers plot, showing the minimum to maximum values. The bar plots show the median with all data points displayed.

### N-acetylglucosamine (*nagB*) and hexuronates (*kdgK*) are context-dependent nutrient sources

The *galK* and *hyb*-deficient SL1344 mutants accentuate context-dependency in carbohydrate and energy metabolism. Similarly, *nagB* and *kdgK* provide additional examples, as their attenuated fitness is observed only in a subset of mouse models. *nagB* encodes glucosamine-6-phosphate deaminase, important in N-acetylglucosamine degradation. A *nagB*-deficient SL1344 mutant shows a 5-fold attenuation in the germ-free model at day 2 post-infection, but a competitive index close to 1 in streptomycin-pretreated models. In LCM models, the *nagB* mutant shows a 50-fold attenuation, significantly more than in the OligoMM^12^ model and any other model (**Fig. 2F**). This highlights and confirms the specific importance of NagB in the LCM model (Nguyen et al., 2024). *kdgK* encodes 2-dehydro-3-deoxygluconokinase, involved in β-glucuronide (hexuronates) degradation. Post-antibiotic stress induces the inducible nitric oxide synthase (iNOS) in the cecal mucosa, leading to monosaccharide oxidation to sugar acids (Faber et al., 2016). KdgK catalyzes the penultimate step in D-fructuronate, D-mannonate, and D-glucuronate degradation (**Fig. 2G**). The *kdgK*-deficient SL1344 mutant is attenuated only in streptomycin-pretreated and germ-free models, showing a wild-type-like competitive index in LCM and OligoMM^12^ models (**Fig. 2H**). This suggests that post-antibiotic stress creates a niche for *S*. Typhimurium colonization, particularly hexuronates of D-fructose and D-mannose. These observations are also consistent for Δ*dgoK* (D-galactonate) and Δ*uxaC* (D-galacturonate) mutants, which show consistently lower competitive indices in streptomycin-pretreated models than in gnotobiotic models (**Fig. S2A-B**). However, this pattern appears to be specific, as not every hexuronate displays it. Specifically, Δ*idnD* (L-idonate) shows no competitive disadvantage in all mouse models (**Fig. S2C**). Thus, Δ*nagB* and Δ*kdgK* mutants, as representative of hexuronates, further highlight the context-dependent nature of carbohydrate metabolism across mouse models.

### D-fructose provides a niche important in intraspecies competition

To identify carbohydrates that are important for *S*. Typhimurium intraspecies competition, we analyzed the SL1344 mutant pool in the presence of a niche competitor, *S*. Typhimurium strain ATCC14028S. Using the streptomycin-pretreated 129S6/SvEvTac mouse model, SL1344 was competed with ATCC14028S at ratios of 1:50 and 1:1000. Despite ATCC14028S’s higher abundance, it did not alter SL1344 loads in fecal and cecal samples, consistent with previous findings (Gül et al., 2023). SL1344 loads exceeded 10^10^ CFU per gram of feces by day 1 post-infection (**Fig. 2I**). Most samples had Shannon evenness scores above 0.9, allowing robust fitness evaluation (**Fig. 2K**). ATCC14028S only had a significant effect on three mutants: *pgl*, *pfkA*, and *fruK*, with *fruK* being roughly 100-fold attenuated at day 2 post infection (**Fig. 2L, Fig. S3A-B**). These findings indicate that carbohydrate competition is highly specific and potentially centered on D-fructose utilization for *Salmonella serovars*, with most SL1344 mutants showing no significant change in fitness in the presence of ATCC14028S. In **Table S7**, the competitive indices for all metabolic mutants in intraspecies competition are listed.

### Hexoses provide the basis for successful colonization

Our previous experiments highlighted three key metabolic genes: *fruK*, *galK*, and *manA*, involved in the degradation of D-fructose, D-galactose, and D-mannose. A D-glucose-associated mutant was initially absent in the SL1344 pool due to the lack of an ideal cytosolic enzyme. D-glucose can be transported by the primary transporter PtsG, as well as by several other transporters (Mayer & Boos, 2005). In the streptomycin-pretreated C57BL/6J model, the *ptsG*-deficient SL1344 mutant showed a 5-10-fold fitness defect from day 2 post-infection (**Fig. 3A**). However, this differed in the gnotobiotic LCM mouse model, where a *ptsG* mutant did not display a significant colonization defect (Nguyen et al., 2024). Integrating *manA*, *fruK*, and *galK* mutations with *ptsG* deficiency aimed to further assess the importance of D-glucose utilization. Multimutants were constructed systematically, starting with the most attenuated mutant, *manA*, followed by *fruK*, *galK*, and finally *ptsG*.

**Figure 3.**
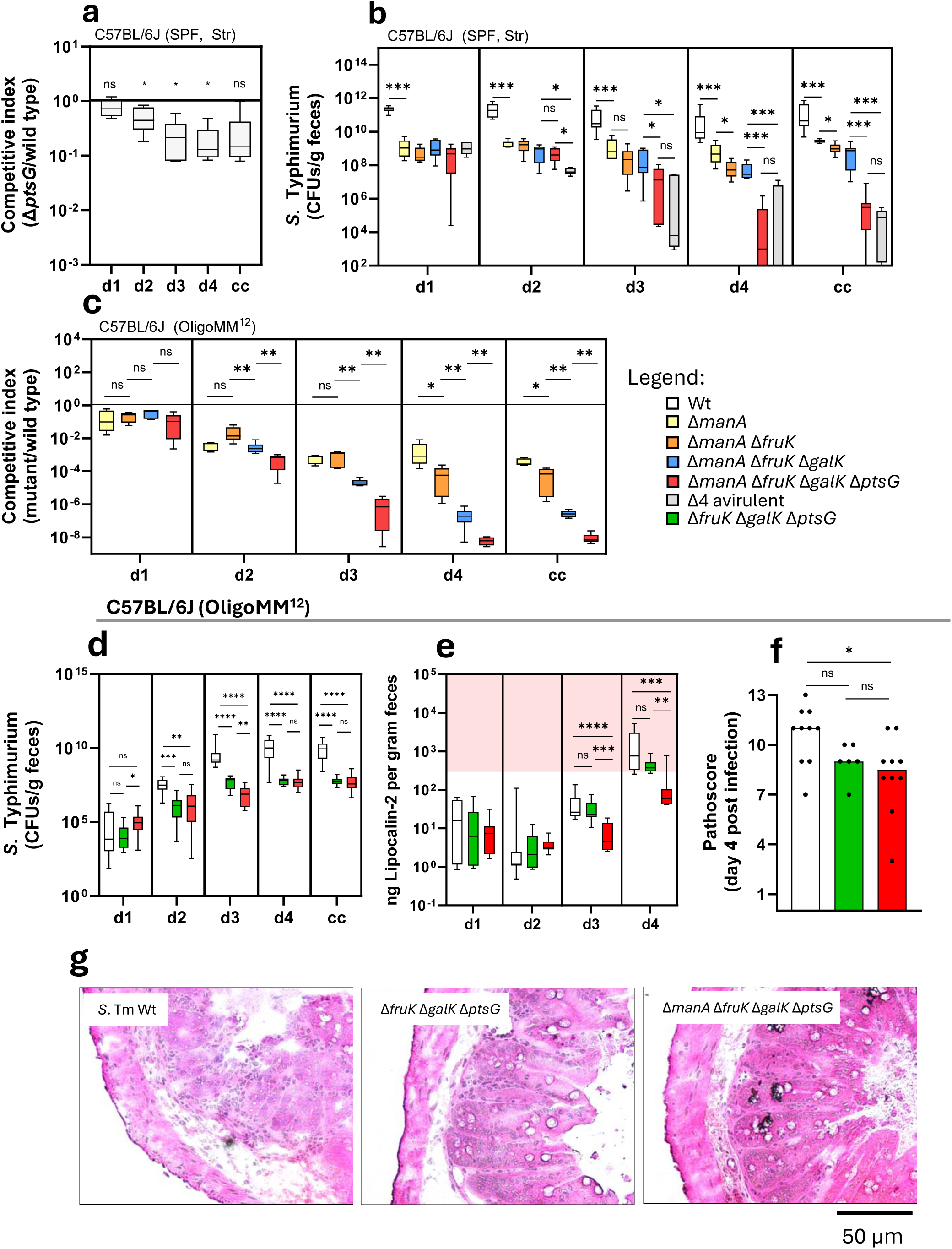
Hexoses provide the basis for *Salmonella* colonization. **A** The competitive experiment was performed in streptomycin-pretreated C57BL/6J mice, lasting 4 days post-infection, comparing the SL1344 wild type with a *ptsG* mutant, which lacks the main glucose transporter (mice n=6, at least two independent experiments). **B** Single infection of streptomycin pretreated C57BL/6J mice with either SL1344 wild type (mice n=10), Δ*manA* (mice n=6), Δ*manA* Δ*fruK* (mice n=6), Δ*manA* Δ*fruK* Δ*galK* (mice n=7), and Δ*manA* Δ*fruK* Δ*galK* Δ*ptsG* quadruple mutant (mice n=7), or an avirulent (Δ*invG* and Δ*ssaV*) quadruple mutant (mice n=5, at least two independent experiments). The bacterial loads are plotted in CFUs per gram feces and cecum content for day 1 to 4 post infection. **C** The competitive experiment of C57BL/6J mice associated with the OligoMM^12^ microbiota with SL1344 wild type (mice n=23) and either a Δ*manA* (mice n=5), Δ*manA* Δ*fruK* (mice n=6), Δ*manA* Δ*fruK* Δ*galK* (mice n=6), or Δ*manA* Δ*fruK* Δ*galK* Δ*ptsG* quadruple mutant (mice n=7, at least two independent experiments). The competitive index for each mutant is plotted and statistically compared to the SL1344 wild type. **D** Single infection of C57BL/6J mice associated with the OligoMM^12^ microbiota with SL1344 wild type (mice n=12), Δ*fruK* Δ*galK* Δ*ptsG* (mice n=10), and Δ*manA* Δ*fruK* Δ*galK* Δ*ptsG* mutant (mice n=11, at least two independent experiments). The bacterial loads are plotted in CFUs per gram feces and cecum content. **E** Lipocalin-2 ELISA as marker for gut inflammation for the OligoMM^12^ model infected with either SL1344 wild type, Δ*fruK* Δ*galK* Δ*ptsG,* and Δ*manA* Δ*fruK* Δ*galK* Δ*ptsG* mutant. The red color indicates an inflammatory state starting at 5×10^2^ ng lipocalin-2 per gram of feces **F** Histopathology scoring of the cecum tissue of OligoMM^12^ mice at day 4 post infection. **G** Microscopy images of a representative part of the cecum tissue from OligoMM^12^ mice infected with SL1344, wild type, Δ*fruK* Δ*galK* Δ*ptsG,* and Δ*manA* Δ*fruK* Δ*galK* Δ*ptsG*, as indicated above the picture. All competitive experiments are presented in a box-and-whiskers plot, showing the minimum to maximum values. The bar plots show the median with all data points displayed.

To evaluate their colonization capacity, SL1344 multimutants were tested in the streptomycin-pretreated C57BL/6J mouse model in a single infection experiment (**Fig. 3B**). The SL1344 wild type reached ≈10^10^ CFU/g feces by day 1 post-infection, maintaining this density until day 4. The *manA*-deficient strain grew significantly lower day 1, staying roughly 100-fold lower than the wild type. The double (Δ*manA* Δ*fruK*) and triple (Δ*manA* Δ*fruK* Δ*galK*) mutants showed stable colonization over 4 days. However, while the quadruple mutant (Δ*manA* Δ*fruK* Δ*galK* Δ*ptsG*) had similar loads to the triple mutant initially, densities collapsed by days 3-4 (**Fig. 3B**). Previous work showed gut microbiota regrowth by days 3-4 if *Salmonella* mutants fail to trigger gut inflammation (Coombes et al., 2005; Hapfelmeier et al., 2005; Stecher et al., 2007). To test this, we used an avirulent isogenic mutant with disrupted T3SS-1 and −2 (Δ*invG* Δ*ssaV* Δ*manA* Δ*fruK* Δ*galK* Δ*ptsG*) showed similar colonization kinetics to the quadruple mutant, indicating that the quadruple mutant is incapable of triggering gut inflammation (**Fig. 4B**).

**Figure 4:**
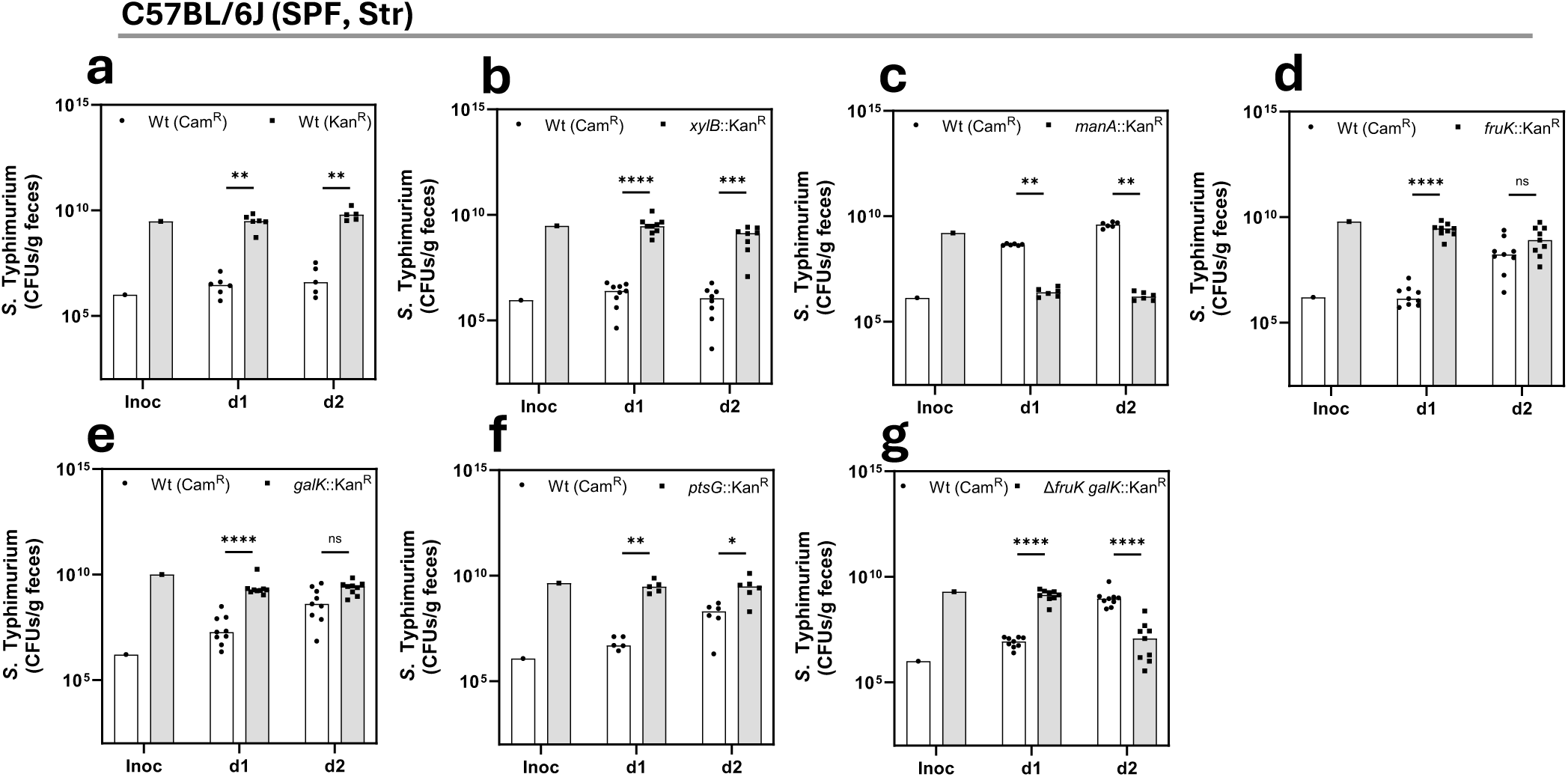
Hexoses provide an exploitable niche for *S*. Typhimurium. Competitive experiments were performed in the streptomycin-pretreated C57BL/6J model, where the carbohydrate utilization mutant is in large excess and the SL1344 wild type constitutes only 0.1% of the infection mix. The bacterial loads for each strain were determined by differential plating on MacConkey agar plates with kanamycin and chloramphenicol antibiotics. **A-G** The individual bacterial loads are plotted in CFUs per gram of feces, including the inoculum, with the strains indicated at the top of the respective diagram for the 2-day infection period (minimum number of mice n=6, at least two independent experiments). The bar plots show the median with all data points displayed.

To assess the role of gut microbiota in the competitive fitness of carbohydrate utilization mutants, we tested the quadruple (Δ*manA* Δ*fruK* Δ*galK* Δ*ptsG*) mutant in the OligoMM^12^ model against the SL1344 wild type. The single (Δ*manA*), double (Δ*manA* Δ*fruK*), and triple (Δ*manA* Δ*fruK* Δ*galK*) mutants were also tested. On day 1 post-infection, competitive indices were similar for all mutants (**Fig. 3C**). However, by days 3 and 4, distinct attenuation levels appeared, with a gradual decrease in fitness from single to quadruple mutants. By day 4, the quadruple mutant’s bacterial loads in fecal and cecal samples were below the detection limit (**Fig. S4**). This highlights the importance of D-glucose, D-fructose, D-mannose, and D-galactose utilization for successful *S*. Typhimurium colonization. Although the single mutant *galK* was initially not attenuated in the OligoMM^12^ model (**Fig. 2E**), its addition to the multi-mutant caused a significant fitness loss compared to the double mutant (**Fig. 3C**). This was probably due to nutritional redundancy in the single *galK* mutant, suggesting that D-galactose is also a context-independent nutrient source.

### The quadruple sugar mutant induces less inflammation in the OligoMM^12^ model

*S*. Typhimurium divides the labor of virulence expression into two phenotypically different subpopulations: virulent and avirulent, due to the high metabolic cost of virulence gene expression (Ackermann et al., 2008; Arnoldini et al., 2014; Diard et al., 2013). To further assess competition against the OligoMM^12^ microbiota and the ability to cause inflammation, we performed single infections with wild-type SL1344, the quadruple mutant (Δ*manA* Δ*fruK* Δ*galK* Δ*ptsG*), and a triple mutant (Δ*fruK* Δ*galK* Δ*ptsG*), which retains a functional *manA* gene. The wild type reached ≈10^9^ CFU/g feces by day 3, while both mutants showed 100-fold lower colonization by days 2-3 (**Fig. 3D**). The triple mutant reached ≈10^6^ CFU/g feces faster than the quadruple mutant, but both strains reached this density by day 4. Unlike in the streptomycin-pretreated C57BL/6J model, the quadruple mutant stably colonized the gut without being displaced by regrowing microbiota, although at significantly lower levels than the wild type (**Fig. 3D**). These data show that while both mutants can stably colonize without wild-type competition, their population size is about 100-fold smaller. To evaluate inflammation, we measured lipocalin-2 in feces and analyzed cecum histopathology (**Fig. 3E-F**). Infection with both the wild type and the triple mutant resulted in high lipocalin-2 levels and severe enteropathy by day 4, indicating pronounced mucosal inflammation (**Fig. 3E-G**). Lower levels of inflammation were observed in mice infected with the quadruple mutant, with both lipocalin-2 and enteropathy (less edema) being significantly lower compared to the wild type (**Fig. 3E-G**). In conclusion, D-mannose (Δ*manA*), D-fructose (Δ*fruK*), D-galactose (Δ*galK*), and D-glucose (Δ*ptsG*) are essential nutrient sources for efficient colonization by *S*. Typhimurium and inducing inflammation during the 4-day infection period in the OligoMM^12^ model.

### *S.* Typhimurium exploits hexoses during colonization

The previous competition experiments focused on mutant fitness in classic 1:1 competitive experiments. However, by supplying the carbohydrate mutant in large excess compared to the wild type (0.1% of the infection mixture), we can determine if the wild type can efficiently utilize the niche that is opened up by the mutant to reach the same level within days, as previously shown for D-galactitol utilization (Gül et al., 2023). This experimental change shifts the focus from the mutant to the wild type, excluding pleiotropic effects that can occur in metabolic mutants (Boulanger et al., 2022; Boulanger et al., 2021). We performed these experiments with various carbohydrate utilization mutants in the streptomycin-pretreated C57BL/6J model. Two identical SL1344 wild-type strains with different antibiotic resistances maintained a stable gut density and ratio between each other (**Fig. 4A**). Using the *xylB* mutant (xylulokinase, D-xylose), which showed no change in competitive index (**Table S2-S6**), confirmed this (**Fig. 4B**). When there is no change in exploitable metabolic capacity, both strains will stay at the same density and ratio. The *manA*-deficient strain was displaced by the wild type by day 1 post-infection, consistent with its severe colonization defect in this mouse model (**Fig. 4C**). For *fruK* and *galK* mutants, the wild type caught up by day 2 (**Figs. 4D-E**). D-galactose provided a more favorable niche than D-fructose, indicated by wild type catch-up kinetics (1.9 × 10^7^ vs. 1.4 × 10^6^ CFU/g feces, respectively. The wild type also caught up with the *ptsG* mutant, though more slowly than with the *galK* and *fruK* mutant (**Fig. 4F**).

To further assess the roles of D-fructose and D-galactose utilization, we tested a *fruK galK* double mutant. The wild type not only caught up faster but displaced the double mutant within 2 days post-infection, a more pronounced phenotype than the single *fruK* or *galK* mutants (**Fig. 4G compared to D and E**). In conclusion, D-fructose, D-galactose, and D-glucose provide exploitable niches for *S*. Typhimurium. For D-mannose, it remains unclear if displacement of the mutant or metabolic exploitation was decisive for the catch up of the wild type. D-galactose is more favorable than D-fructose in the streptomycin-pretreated C57BL/6J model, even though the Δ*fruK* mutant’s fitness was more attenuated than that of the Δ*galK* mutant (**Fig. 2B**). The role of glucose utilization is tougher to discern since the *ptsG* mutation only impairs one of the possible D-glucose transporters. However, inactivating PtsG creates a metabolic gap large enough to be exploited by the wild type.

### Free monosaccharides are abundant in cecum contents of mice

Our experiments show that genes associated with the utilization of D-mannose (*manA*), D-fructose (*fruK*), D-glucose (*ptsG*), and D-galactose (*galK*) are context-independent in our five mouse models of *S*. Typhimurium colonization. N-acetylglucosamine (*nagB*, GlcNAc) and hexuronates (*kdgK*, β-glucuronide, *dgoK*, D-galactonate; *uxaC*, D-galacturonate) serve as context-dependent niches in certain mouse models. To determine carbohydrate availability in the cecum, we collected contents from gnotobiotic OligoMM^12^ and germ-free mice, and analyzed them using LC-MS. This was compared with previously published data from C57BL/6J SPF mice (**Fig. 5A**) (Nguyen et al., 2024). The hexoses D-glucose, D-galactose, and D-mannose are consistently lower in the C57BL/6J SPF mice than in the germ-free and OligoMM^12^ models. Surprisingly, D-glucose is similarly abundant in the OligoMM^12^ model as in germ-free mice, indicating that the OligoMM^12^ consortium does not specifically limit this essential nutrient (**Fig. S5A**). A similar pattern is observed for amino sugars, which are generally lowest in C57BL/6J SPF mice and the OligoMM^12^ model, reflecting competition for amino sugars and hexoses among the resident microbiota (**Fig. S5B**). This pattern also applies to the pentose D-ribose. L-arabinose has the highest abundance in the OligoMM^12^ model, indicating the presence of microbiota that can liberate L-arabinose from arabinan polymers without consuming the freed monomer. D-xylose shows similarly high abundance in both the OligoMM^12^ and SPF C57BL/6J mice, compared to germ-free animals (**Fig. S5C**). Regarding deoxy sugars, L-rhamnose and L-fucose were previously shown to be of microbial origin (Meier et al., 2023). These sugars can be utilized by some microbiota members for fermentation to produce the electron acceptor 1,2-propanediol (Faber et al., 2017) (**Fig. S5D**). Specifically, L-rhamnose is abundant in the OligoMM^12^ model compared to the other two, suggesting that L-rhamnose is produced but not efficiently consumed by the microbiota (**Fig. S5D**). Hexuronates, such as D-glucuronic acid and D-mannuronic acid, are generally most abundant in germ-free mice, except for D-galacturonic acid. This explains the attenuated fitness of the hexuronate-utilizing *S*. Typhimurium *kdgK* mutant in this particular mouse model (**Fig. S5E**). In conclusion, D-glucose and D-galactose were found in the millimolar range, while D-mannose, N-acetylglucosamine, and D-glucuronic acid were in the micromolar range (**Fig. 5A**). The C57BL/6J SPF model consistently had lower free monosaccharide levels than the OligoMM^12^ model, most likely due to consumption by the complex microbiome, reinforcing the essentiality of a complex microbiome for metabolic exploitation.

**Figure 5:**
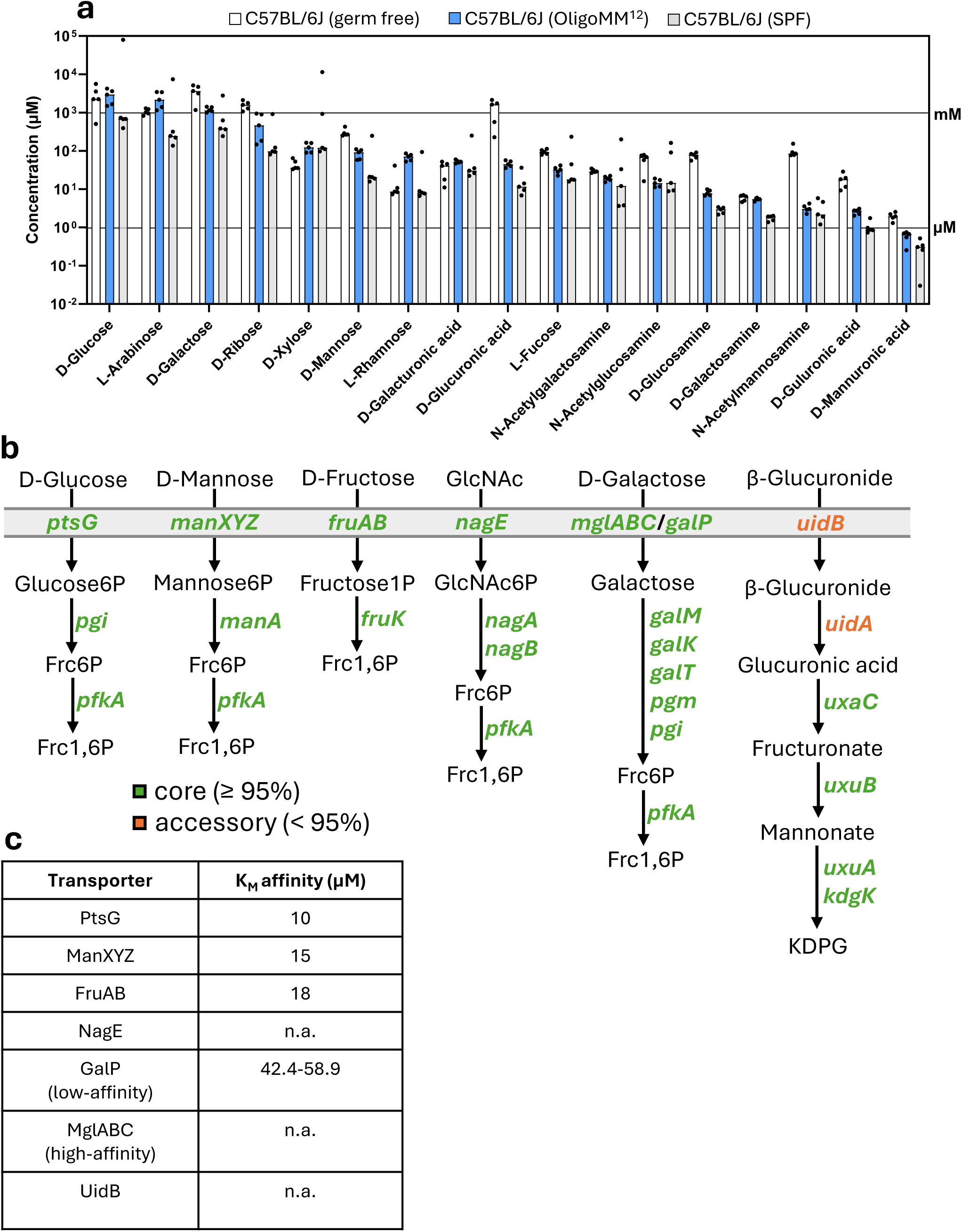
Free monosaccharides are abundant in cecum contents of mice. **A** Absolute measurement of free monosaccharides in cecum contents of C57BL/6J mice associated with the OligoMM^12^ microbiota (mice n=5) and germ free (mice n=5) quantified by LC-MS. For comparison a previously published dataset of C57BL/6J with a complex specific pathogen free (SPF) microbiota was included (mice n=5) (Nguyen et al., 2024). The black lines indicate mM and µM concentration, respectively. **B** A schematic representation of the D-glucose, D-mannose, D-fructose, N-acetylglucosamine (GlcNAc), D-galactose, and β-glucuronide (hexuronate) degradation pathways. If a gene is present in at least 95% of all analyzed genomes, it is termed core (green). Less than 95%, it is termed accessory (orange). The full names of the abbreviations are listed in **Table S12**. **C** Bacterial transporters with the respective K_M_ affinity are shown in µM. PtsG and ManXYZ from *S*. Typhimurium, (Stock et al., 1982); FruAB from *P. aeruginosa*, (Durham & Phibbs, 1982); GalP from *E. coli*, (McDonald et al., 1997). K_M_ affinities for NagE, MglABC, and UidB are not available (n.a.).

Based on these considerations, we hypothesized that genes for utilizing critical carbohydrates fuelling pathogen growth in the host’s gut lumen should be universally present among *S*. *enterica* strains. To understand the distribution of these metabolic systems within *Enterobacteriaceae*, we analyzed the presence of genes involved in carbohydrate utilization across four genera: *Salmonella*, *Escherichia*, *Shigella*, and *Citrobacter*. D-glucose, D-mannose, D-fructose, N-acetylglucosamine, and D-galactose utilization genes are retained in over 95% of genomes analyzed **(Fig. 5B)**. Only the transport and the initial step of β-glucuronide degradation are linked to the accessory genome, which are not found in *Salmonella* (**, Table S8**). With the exception of D-galactose and hexuronates, D-glucose, D-mannose, D-fructose, and GlcNAc are transported by the phosphotransferase system (PTS) (Deutscher et al., 2014; Görke & Stülke, 2008). PTS transporters have high affinity for sugars in the low micromolar range, while GalP, a low-affinity transporter, has approximately 5-fold lower affinity (**Fig. 5C**). In contrast, human glucose transporters have affinities ranging between 0.2 and 17 mM (Zhao & Keating, 2007). This suggests that enteric bacteria have evolved to efficiently utilize monosaccharides in low micromolar concentrations, consuming what is not utilized by the host.

## Discussion

To assess the metabolic pathways that promote enteropathogen growth in the gut, we screened a mutant pool lacking key enzymes involved in carbohydrate utilization pathways of *S*. Typhimurium across five different mouse models. This approach establishes a fast and cost-effective method to determine the metabolic requirements of *S*. Typhimurium for colonizing different environments. This confirmed the critical role of H_2_/fumarate respiration in the LCM gut (Maier et al., 2013; Nguyen et al., 2020) and emphasized the importance of fumarate respiration for *Enterobacteriaceae* (Jones et al., 2011; Nguyen et al., 2020; Schubert & Unden, 2022; Schubert & Unden, 2023; Schubert et al., 2021; Yoo et al., 2024) in a wide range of different mouse models. Here we further identified metabolic genes associated with the utilization of D-glucose (*ptsG*), D-mannose (*manA*), D-fructose (*fruK*), and D-galactose (*galK*) to be context-independent, while N-acetylglucosamine (*nagB*) and hexuronates (*kdgK*, *dgoK*, and *uxaC*) are context-dependent for *S*. Typhimurium colonization across various mouse models. Combining mutations of context-independent metabolic genes (Δ*manA* Δ*fruK* Δ*galK* Δ*ptsG*) in one SL1344 strain progressively attenuated gut-luminal growth, underscoring their significance as the basic metabolic requirements for successful *S*. Typhimurium colonization.

D-glucose, D-mannose, D-galactose, and D-fructose are prevalent in dietary plants, host glycans, and various other sources (Porter & Martens, 2017). The monomer composition of different foods predominantly consists of D-glucose, reflecting its natural abundance (Castillo et al., 2022; Englyst et al., 2007; Larke et al., 2023). Quantifying these carbohydrates in the cecal contents of gnotobiotic OligoMM^12^ and germ-free mice revealed D-glucose and D-galactose in the millimolar range, while others, including N-acetylglucosamine and D-glucuronic acid, were in the micromolar range. These concentrations exceed bacterial transport affinities by one or two orders of magnitude (**Fig. 5C**). Interestingly, unperturbed SPF C57BL/6J mice had consistently lower concentrations of D-glucose, D-mannose, and D-galactose, compared to the OligoMM^12^ model, highlighting the importance of a complex microbiome for efficient metabolic exploitation to provide colonization resistance. Even though monosaccharide concentrations are still considerable in SPF C57BL/6J mice, *S*. Typhimurium has substantial difficulty colonizing an unperturbed C57BL/6J SPF mouse model (Nguyen et al., 2020). This suggests that either these monosaccharide concentrations are too low or that metabolic exploitation of monosaccharides is not the only key factor in preventing *S*. Typhimurium colonization. D-fructose was shown to be available in SPF C57BL/6 mice at concentrations that support *Enterococcus* colonization (Isaac et al., 2022). Furthermore, D-fructose seems to be an important nutrient niche for intraspecies competition between two *Salmonella serovars*. Glycan-degraders like *Bacteroides thetaiotaomicron* release monosaccharides from complex polysaccharides, making them accessible to other microbiota members or pathogens (Ng et al., 2013). Free monosaccharide concentrations in germ-free mice often times exceeded those in the OligoMM^12^ measurements, such as in the case of D-galactose and D-mannose (**Fig. 5A**). This suggests that a sterile host does not efficiently consume monosaccharides, highlighting the importance of the synergy between host and microbiota in limiting these nutrients.

Previous studies showed that pathogenic *E. coli* (EDL933) utilizes D-galactose, hexuronates, D-mannose, and D-ribose, while commensal *E. coli* (MG1655) relies on D-gluconate and N-acetylneuraminic acid (Chang et al., 2004; Doranga et al., 2024; Fabich et al., 2008). This emphasizes the fact that *S*. Typhimurium and *E. coli* have a highly similar metabolic resource overlap, underlining the observation that *E. coli* is a key competitor against pathogenic *S*. Typhimurium in the mammalian gut (Spragge et al., 2023; Velazquez et al., 2019). In conclusion, *E. coli* and *Salmonella* spp. prefer monosaccharides to support their growth in the mammalian gut lumen, as evidenced by the presence of the necessary metabolic enzymes in both genera (**Fig. 5B**). The availability of free monosaccharides in the mouse gut is sufficient for microbial utilization, highlighting the critical role of metabolic exploitation by the microbiota in limiting pathogen invasion. Furthermore, phylogenetic analysis suggests that these metabolic systems are conserved across *Salmonella*, *Escherichia*, *Citrobacter*, and *Shigella*, indicating that these metabolic strategies presented here are common among *Enterobacteriaceae*.

In humans, two primary glucose transporters exist: the passive GLUT transporter and the sodium-dependent SGLT transporters, with a Michaelis-Menten constant (K_M_) between 0.2 and 17 mM (Zhao & Keating, 2007). K_M_ essentially refers to the amount of substrate required for an enzyme to operate at half its maximal velocity, wherein the substrate affinity can be markedly lower. This K_M_ is an order of magnitude lower than that of bacterial hexose transporters, which can uptake monosaccharides potentially in the nanomolar range (**Fig. 5C**). Thus, even if free monosaccharide levels are lower than those in the OligoMM^12^ model or SPF C57BL/6J mice (**Fig. 5A**), they can still support some level of *S*. Typhimurium colonization. In conclusion, we provided fitness information for 35 metabolic genes across five different mouse models to differentiate nutrient sources into context-independent and context-dependent. This information will guide the future design of probiotics and microbiota consortia to actively reduce these metabolites to limit nontyphoidal *Salmonella* infection.

## Acknowledgements

We would like to acknowledge and thank the staff at the ETH animal facilities (EPIC and RCHCI; especially Manuela Graf, Katharina Holzinger, Dennis Mollenhauer, Sven Nowok, and Dominik Bacovcin), and extend many thanks to members of the Hardt, Sunagawa, Vorholt, and Slack labs, as well as the NCCR Microbiomes, for their helpful comments and discussions. Many thanks to Yassine Cherrak, Leanid Laganenka, and Gottfried Unden for providing helpful comments.

## Author contributions

C.S. and W.-D.H. conceived and designed the experiments. C.S. performed the *in vivo* experiments. B.N., A.S., and J.N. collected and analyzed monosaccharides by LC-MS. A.S. and L.F. provided the bioinformatics pipeline for WISH analysis. B.D. and Y.S. provided support for WISH barcoding. N.N. performed the bioinformatic analysis for the presence of metabolic systems. C.S. wrote the manuscript with contributions from all authors.

## Methods

### Animals

We used male and female mice aged 8-12 weeks, randomly assigning animals of either sex to the experimental groups. All animal models received the same standard SPF chow in our mouse facility. The mice originated from C57BL/6J or 129S6SvEv/Tac breeders initially obtained from Jackson Laboratories. Mice with a normal complex microbiota were specific pathogen-free (SPF) and bred under full barrier conditions in individually ventilated cage systems at the EPIC mouse facility of ETH Zurich, Switzerland. LCM mice (Maier et al., 2013; Stecher et al., 2010) and OligoMM^12^ mice (Brugiroux et al., 2016) are ex-germ-free C57BL/6J animals that have been associated with either strains of the altered Schädler flora or a genome-guided selection of a representative group of 12 microbiota strains. They are bred in flexible film isolators under strict exclusion of microbial contamination at the isolator facility of the EPIC mouse facility. Germ-free C57BL/6J mice were bred in flexible film isolators under strict exclusion of microbial contamination at the isolator facility of the EPIC mouse facility, ETH Zurich, Switzerland. All studies were conducted in compliance with ethical and legal requirements and were reviewed and approved by the Kantonales Veterinäramt Zürich under licenses ZH158/19, ZH108/22 and ZH109/2022.

### Bacteria and culture conditions

All *Salmonella* strains are isogenic to *Salmonella* Typhimurium SB300, a re-isolate of SL1344. They are listed in **Table S9**. All plasmids used in this study are listed in **Table S10**. All strains were routinely grown overnight at 37°C in Lysogeny broth (LB) with agitation. Strains were stored at −80°C in peptone glycerol broth (2% w/v peptone, 5% v/v glycerol (99.7%)). Custom oligonucleotides were synthetized by *Microsynth* and are listed in **Table S11**.

### Homologous recombination by Lambda Red

Single-gene knockout strains were generated using the lambda-red single-step protocol (Datsenko & Wanner, 2000). Primers were designed with an approximately 40 bp overhanging region homologous to the genomic region of interest and 20 bp binding region corresponding to the antibiotic resistance cassette (**Table S10**). PCR amplification was performed using the plasmid pKD4 for kanamycin resistance or the pTWIST plasmids for WISH tags, which include an ampicillin resistance cassette. DreamTaq Master Mix (Thermo Fisher Scientific) was employed, followed by digestion of the template DNA using FastDigest DpnI (Thermo Fisher Scientific). Subsequently, the PCR product was purified using the Qiagen DNA purification kit. SB300 with either the pKD46 or pSIM5 plasmid was cultured for 3 h at 30°C until early exponential phase, followed by induction with L-arabinose (10 mM, Sigma-Aldrich) or 42°C for 20 min, respectively. The cells were washed in ice-cold glycerol (10% v/v) solution and concentrated 100-fold. Ultimately, the PCR product was transformed by electroshock (1.8 V at 5 ms), followed by regeneration in SOC (SOB pre-made mixture, Roth GmbH, and 50 mM glucose) medium for 2 h at 37°C, ultimately plated on selective LB-agar plates. The success of the gene knockout was verified by gel electrophoresis and sanger sequencing (Microsynth AG). Kanamycin resistance cassettes were eliminated via flippase FLP recombination (Cherepanov & Wackernagel, 1995).

### Homologous recombination by P22 phage transduction

P22 phage transduction was conducted by generating P22 phages containing the antibiotic resistance cassette inserted into the gene of interest from the defined single-gene deletion mutant collection of *S.* Typhimurium (Porwollik et al., 2014). The single-gene knockout mutant was incubated overnight with the P22 phage generated from a wild type SB300 background. The culture was treated with chloroform (1% v/v) for 15 min followed by centrifugation and subsequent sterile filtration (0.44 µM pore size). The P22 phages were subsequently incubated with the recipient strain for 15 minutes and then plated on selective LB-agar plates. This was followed by two consecutive overnight streaks on selective LB-agar plates. Finally, the transduced clone was examined for P22 phage contamination using Evans Blue Uranine (EBU) LB-agar plates (0.4% w/v glucose, 0.001% w/v Evans Blue, 0.002% w/v Uranine). All mutations were verified by gel electrophoresis or Sanger sequencing (Microsynth AG), using the corresponding primers (**Table S11**).

### WISH-barcoding of *S.* Typhimurium

WISH barcodes were introduced, as previously described (Daniel et al., 2024). WISH-tags were amplified from pTWIST using DreamTaq Master Mix (Thermo Fisher Scientific) with WISH_int_fwd and WISH_int_rev primers (**Table S11**) and integrated into *S.* Typhimurium SL1344 (strain SB300), using the λ-red system with pSIM5 (Datsenko & Wanner, 2000). Integration was targeted at a fitness-neutral locus between the pseudogenes *malX* and *malY*, as previously described (Grant et al., 2008). Correct integration was confirmed through colony PCR, and WISH-tags were validated by Sanger sequencing (Microsynth AG), using either the WISH_ver_fwd and WISH_ver_rev primers or the WISH_seq_fwd and WISH_seq_rev primers (**Table S11**). Subsequently, P22 phage lysates were prepared from these generated strains to transduce the WISH-tag into SB300 wild types, controls, and carbohydrate utilization mutants.

### Preparation of the *S.* Typhimurium mutant pool

An individual clone was inoculated in selective LB media and grown overnight at 37°C, containing carbenicillin (100 µg/ml) and kanamycin (50 µg/ml), or chloramphenicol (7.5 µg/ml) to select for the WISH-barcode and the respective mutation. The overnight culture was inoculated into a subculture (5% v/v) and grown for 4 hours at 37°C to the late exponential phase in LB media without antibiotics. At this stage, the strains were pooled in equal volumes, followed by centrifugation (4,500 × g, 4°C, 15 min) and a washing step in 1x PBS. The cell pellets were resuspended in peptone glycerol media to 10% of the original volume and aliquoted in 100 µl volumes in cryo tubes. The pools were stored at - 80°C. This method allows for rapid utilization of the mutant pools, as the SL1344 mutant pool stock is subcultured in LB media only 4 hours prior to infection. To investigate the colonization defects of metabolic mutants in various mouse models, we used aliquots from the same SL1344 mutant pool stocks throughout this study.

### Mouse colonization experiments

For the SB300 mutant pool experiments, 8 to 12-week-old mice were inoculated with cells prepared from a 4 h subculture in LB medium without antibiotics sourced from the pre-mixed cryo stocks. For single and competitive infection experiments, 8 to 12-week-old mice were inoculated with *S*. Typhimurium cells prepared from a culture grown overnight in selective LB. The overnight culture was then used to inoculate (5% v/v) a subculture for 4 h in LB media without antibiotics. In both cases, bacteria were washed once with 1xPBS, and mice were infected with bacteria by gavage (5 × 10^7^ CFU in 50 μl). For the competitive catch-up experiments, the diluted wild-type SB300 strain (chloramphenicol resistant) was diluted 1000-fold and then mixed with either another wild type or carbohydrate mutant (kanamycin resistant) in a 50 µl inoculum mix in equivolume. At the end of the experiments, either day 2 or 4 post infection, animals were euthanized by CO_2_ asphyxiation. Fresh fecal pellets and whole cecal content in PBS (500 μl) using a Tissue Lyser device (Qiagen) before plating to determine the total bacterial population size.

### Sample preparation for the WISH barcode counting

Fecal *S*. Typhimurium cells were enriched in 1 ml LB medium with 100 µg/ml carbenicillin (Carl Roth GmbH) to select for WISH-barcoded strains. Bacterial cells were pelleted, the supernatant was discarded, and then stored at −20°C. DNA extraction from thawed pellets was performed using commercial kits (Qiagen Mini DNA kit) according to the manufacturer’s instructions. For PCR amplification of the WISH-barcodes, 2 µl of isolated genomic DNA sample and 100 nmol of each primer (WISH_Illumina_fwd and WISH_Illumina_rev, see **Table S11**) were used in a DreamTaq MasterMix (Thermo Fisher Scientific). The reaction was conducted with the following cycling program: initial denaturation step at (1) 95°C for 3 min followed by (2) 95°C for 30 sec, (3) 55°C for 30 sec, (4) 72°C for 20 sec for (5) 25 cycles, and a terminal extension step at (6) 72°C for 10 min. PCR products were column purified. We indexed the PCR products for Illumina sequencing by performing a second PCR with nested unique dual index primers using the following program: (1) 95°C for 3 min, (2) 95°C for 30 s, (3) 55°C for 30 s, (4) 72°C for 20 sec, (5) repeat steps 2-4 for 10 cycles, (6) 72°C for 3 min. Afterward, we assessed the indexed PCR product using gel electrophoresis (1% w/v agarose, TAE buffer), pooled the indexed samples according to band intensity, and subsequently purified the library via AMPure bead cleanup (Beckman Coulter) before proceeding to Illumina sequencing. Amplicon sequencing was performed by BMKGENE (Münster, Germany). BMKGENE was tasked with sequencing each sample at a 1 G output on the NGS Novaseq platform, utilizing a 150 bp paired end reads program. Subsequently, the reads were demultiplexed and grouped by WISH-tags using mBARq software (Sintsova et al., 2024). Misreads or mutations of up to five bases were assigned to the closest correct WISH-tag sequence. The WISH barcode counts for each mouse in every experiment are available in **Table S2-S7**. These counts were used to calculate the competitive fitness and Shannon evenness score (7 wild types). WISH counts with less than or equal to 10 were excluded from further analysis and defined as the detection limit, as previously established (Daniel et al., 2024). The Shannon evenness score of the inoculum, excluding the SL1344 wild-type dilution series, was 0.99 after 4 hours of enrichment in selective LB media, indicating no general growth defects in any strain within the pool (**Fig. S6A**). In the streptomycin-pretreated C57BL/6J model, wild-type dilutions at 10^-3^ were detectable at high rates (**Fig. S6B**). This trend was consistent across all tested models (data not shown). Based on this reading, the competitive index of mutants that were below the limit of detection were conservatively set to a competitive index of 10^-3^.

### Competitive index calculation

To calculate the for the mutant pool, the values were determined by dividing the number of observed barcode reads at a specific time point (day 1 post-infection to day 4 post-infection or in cecum content) by the number of barcode reads observed in the inoculum. For the calculation of the Competitive Index (CI), the individual strain fitness of each WISH-barcoded mutant were divided by the mean fitness value of the 7 WISH-barcoded wild-type *S*. Typhimurium control strains. To calculate the statistical significance, the metabolic mutants were compared to the SB300 wild type in the control group. The fitness of the competitive infections with SB300 wild type against the carbohydrate utilization single- and multi-mutants was calculated by dividing the colony-forming units (CFU) per gram of feces of the wild type by those of the mutants.

### Free monosaccharide quantification by LC-MS

Following a previously published protocol (Ruhmann et al., 2014), samples containing free monosaccharides (25 µL) were derivatized with 75 µL of 0.1M 1-phenyl-3-methyl-5-pyrazolone (PMP) in 2:1 methanol:ddH_2_O with 0.4 % ammonium hydroxide for 100 minutes at 70°C. Additionally, each sample was spiked with an internal standard of 10 µM ^13^C6-Glucose, ^13^C6-Galactose and ^13^C6-Mannose (mass 186 Da). For quantification, we derivatized a serial dilution of a standard mix containing D-galacturonic acid, D-glucuronic acid, D-mannuronic acid, D-guluronic acid, D-xylose, L-arabinose, D-glucosamine, L-fucose, D-glucose, D-galactose, D-mannose, N-acetyl-D-glucosamine, N-acetyl-D-galactosamine, N-acetyl-D-mannosamine, D-ribose, L-rhamnose and D-galactosamine. Samples and standards were derivatized by incubation at 70°C for 100 minutes. After derivatization, samples were neutralized with HCl followed by chloroform extraction to remove underivatized PMP as described previously.

Following (Xu et al., 2017), PMP-derivatives were measured on a SCIEX qTRAP5500 and an Agilent 1290 Infinity II LC system equipped with a Waters CORTECS UPLC C18 Column, 90 Å, 1.6 µm, 2.1 mm X 50 mm reversed phase column with guard column. The mobile phase consisted of buffer A (10 mM NH_4_Formate in ddH_2_O, 0.1% formic acid) and buffer B (100% acetonitrile, 0.1% formic acid). PMP-derivatives were separated with an initial isocratic flow of 15% Buffer B for 2 minutes, followed by a gradient from 15% to 20% Buffer B over 5 minutes at a constant flow rate of 0.5 ml/min and a column temperature of 50°C. The ESI source settings were 625°C, with curtain gas set to 30 (arbitrary units), collision gas to medium, ion spray voltage 5500 (arbitrary units), temperature to 625°C, Ion source Gas 1 &2 to 90 (arbitrary units). PMP-derivatives were measured by multiple reaction monitoring (MRM) in positive mode with previously optimized transitions and collision energies. For example, a glucose derivative has a Q1 mass of 511 and was fragmented with a collision energy of 35V to yield the quantifier ion of 175Da and the diagnostic fragment of 217 Da. Different PMP-derivatives were identified by their mass and retention in comparison to known standards. Technical variation in sample processing were normalized by the amount of internal standard in each sample. Peak areas of the 175Da fragment were used for quantification using an external standard ranging from 100 pM to 10 µM.

### Haematoxylin and eosin (HE) staining of tissue

Haematoxylin and eosin (HE) staining of cryo-embedded tissues and subsequent pathoscoring for granulocyte infiltration were performed as described previously (Barthel et al., 2003).

### Lipocalin-2 analysis of feces samples

Lipocalin 2 was detected in fecal samples homogenized in 500 μl sterile PBS using an ELISA assay (DuoSet Lipocalin ELISA kit, DY1857, R&D Systems, Minneapolis, MN, USA). Fecal pellets were diluted 1:20, 1:400, or left undiluted, and concentrations were determined using Four-Parametric Logistic Regression curve fitting.

### Distribution of monosaccharide utilization genes in enteric bacteria

The distribution of genes involved in monosaccharide utilization were based on orthologous gene clusters generated from a dataset of 1158 *Enterobacteriaceae* genomes (Cherrak et al., 2024). In short, Cherrak *et al*, created orthologous gene clusters at different sequence identity thresholds using PIRATE (Bayliss et al., 2019). Gene clusters with relevant functional annotations (**see Fig. 5B**) were extracted and distribution patterns were assessed using Python 3.7.6 across four *Enterobacteriaceae* genera: *Citrobacter, Escherichia, Shigella* and non-typhoidal *Salmonella.* Each genus was randomly subsampled to 65 genomes to ensure even distribution. Gene clusters were classified as core if present in at least 95% of the selected genomes. All other gene clusters were classified as accessory.

### Statistical analysis

No statistical methods were used to predetermine sample sizes. The current sample sizes (n ≥ 5) are similar to those reported in previous publications (Stecher et al., 2004). Data collection and analysis were not performed blind to the conditions of the experiments. Only animals with a Shannon evenness score below 0.9, as previously determined as the cutoff (Maier, Diard, et al., 2014), were excluded from the analyses; otherwise, no animals were excluded. Where applicable, the two-tailed Mann-Whitney U test was employed to assess statistical significance, as specified in the figure legends. Statistical analyses were performed using GraphPad Prism 9 for Windows. P values of p>0.05 were considered not significant (ns), while p<0.05 was denoted as (*), p<0.01 as (**), p<0.001 as (***), and p<0.0001 as (****).

## Supplemental Figures

**Figure S1.**
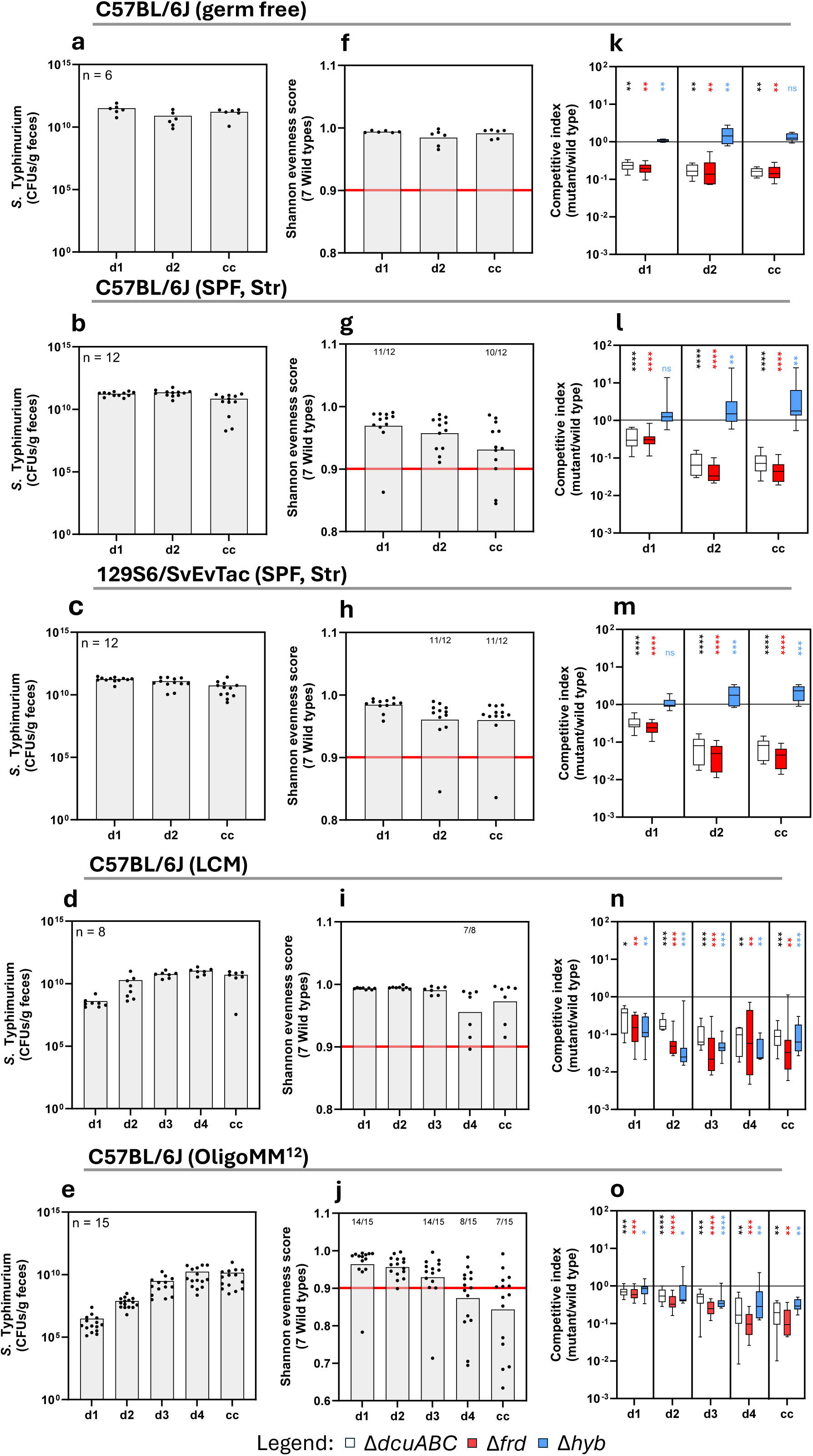
WISH-barcoded control strains provide reproducible data throughout all tested mouse models. The overview shows the *S*. Typhimurium bacterial loads for the mutant pool, Shannon evenness score of the 7 wild types, and the competitive indices of the three control mutants, Δ*dcuABC*, Δ*frd*, and Δ*hyb* in all tested mouse models. The mouse models are indicated above the respective results. **A-E** Black dots indicate total S. Typhimurium colony-forming units per gram of feces and cecum content (CFUs/g), and the sample size for each mouse model is noted. **F-J** Shannon evenness score (SES) was calculated for the 7 WISH-barcoded SL1344 wild types. The red line indicates the SES of 0.9, which was the cutoff for further analysis. The number above the bar indicates how many samples are within this threshold. **K-O** The competitive indices (CI) of the three metabolic mutants used as controls in the *S*. Typhimurium mutant pool, Δ*dcuABC*, Δ*frd*, and Δ*hyb*, was statistically compared to the SB300 wild type in the control group. The black line indicates a wild-type CI at 1. All competitive experiments are presented in a box-and-whiskers plot, showing the minimum to maximum values. The bar plots show the median with all data points displayed.

**Figure S2.**
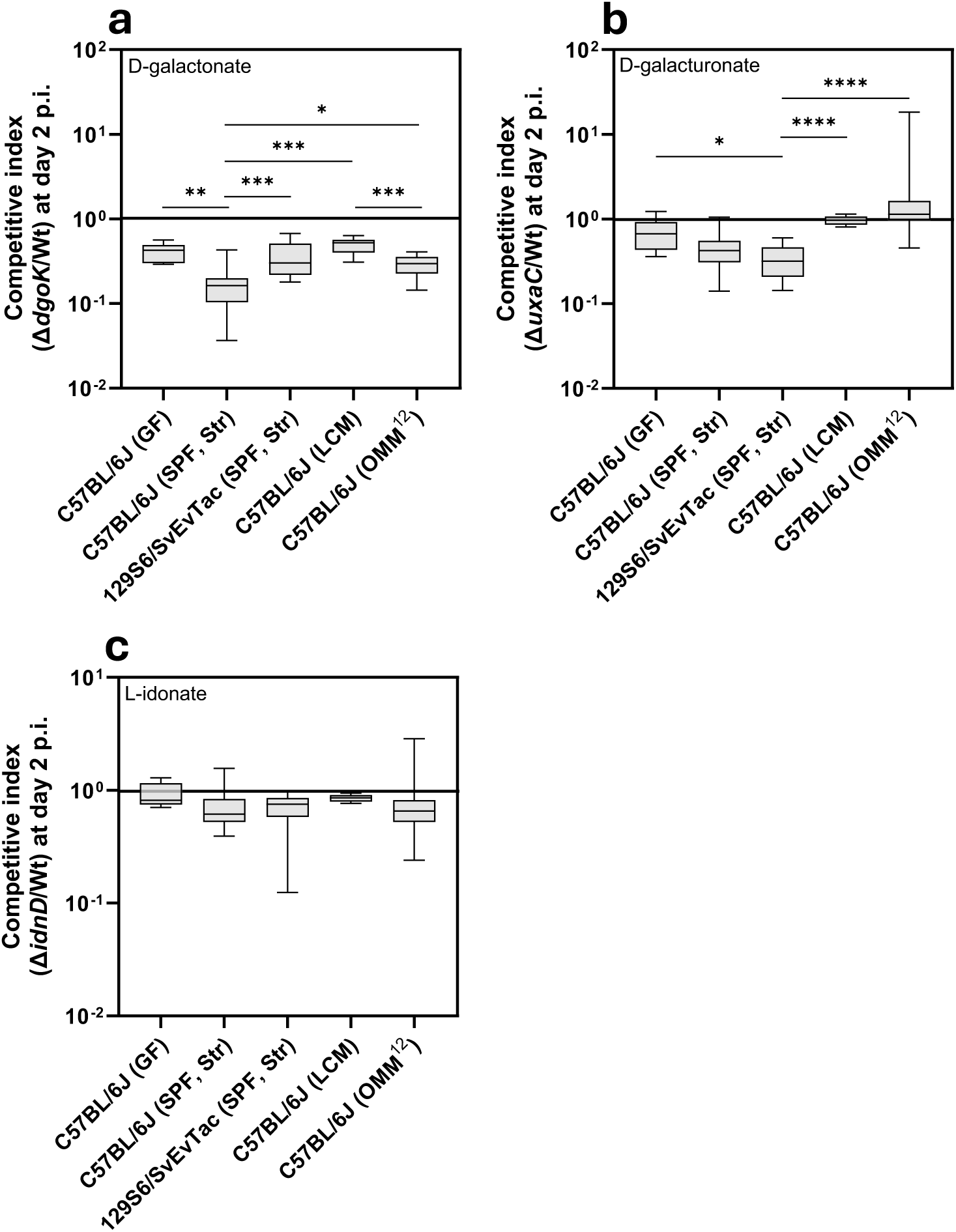
D-galactonate, D-galacturonate, and L-idonate are utilized in a context-dependent manner. **A-C** Competitive indices 2 days post infection for Δ*dgoK* (D-galactonate), Δ*uxaC* (D-galacturonate), and Δ*idnD* (L-idonate) mutants for all five mouse models, as indicated on the x-axis. All competitive experiments are presented in a box-and-whiskers plot, showing the minimum to maximum values.

**Figure S3:**
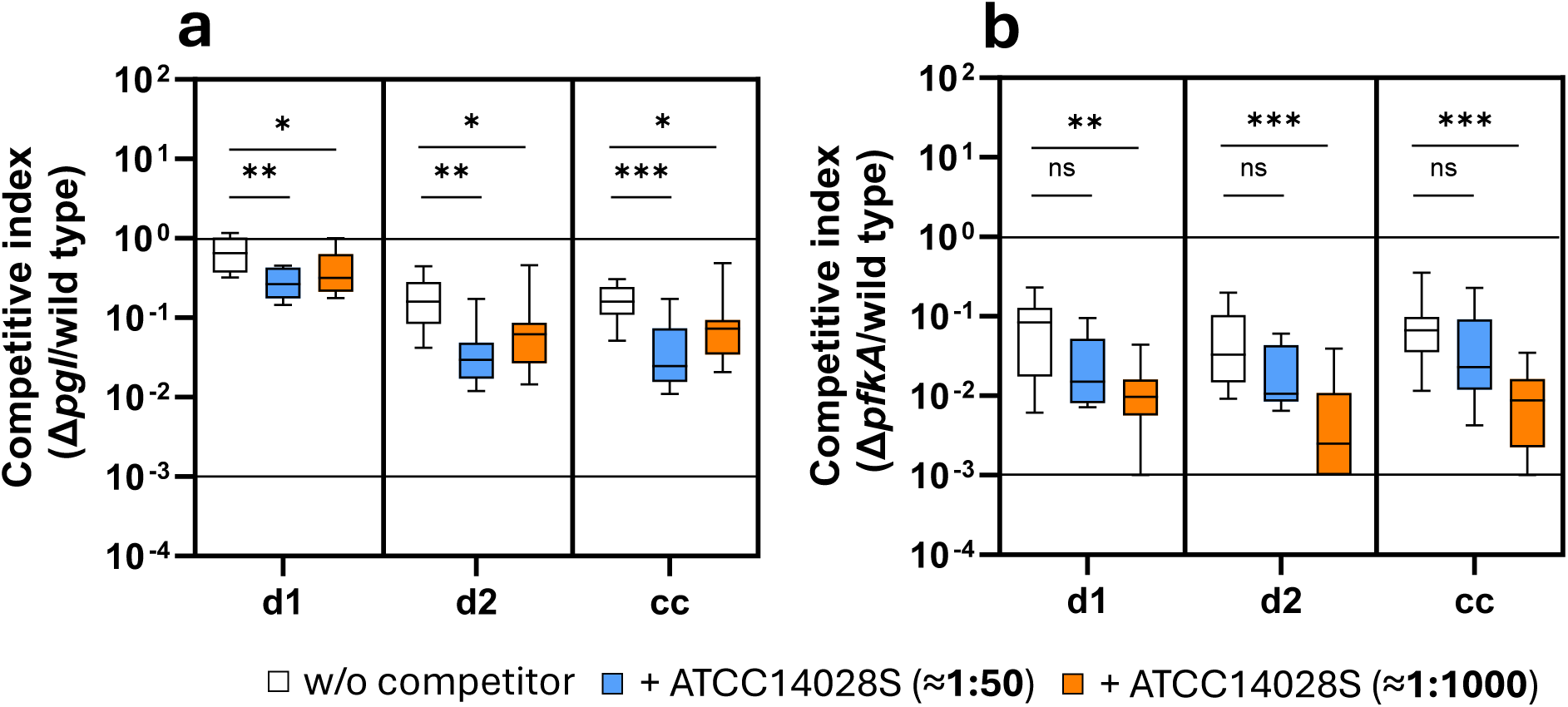
Intraspecies competition between SL1344 and ATCC14028S is highly specific. The WISH-barcoded SL1344 mutant pool was competed with *Salmonella* Typhimurium ATCC14028S in a 1:50 (mice n=9) and 1:1000 ratio (mice n=9, at least two independent experiments). **A-B** The competitive index of the SL1344 *pgl* and *pfkA* mutant is shown, in absence and presence of the competitor ATCC14028S at both ratios. *pgl* encodes the enzyme for the second step of the oxidative pentose phosphate pathway, while *pfkA* encodes phosphofructokinase, a key enzyme in glycolysis. All competitive experiments are presented in a box-and-whiskers plot, showing the minimum to maximum values.

**Figure S4:**
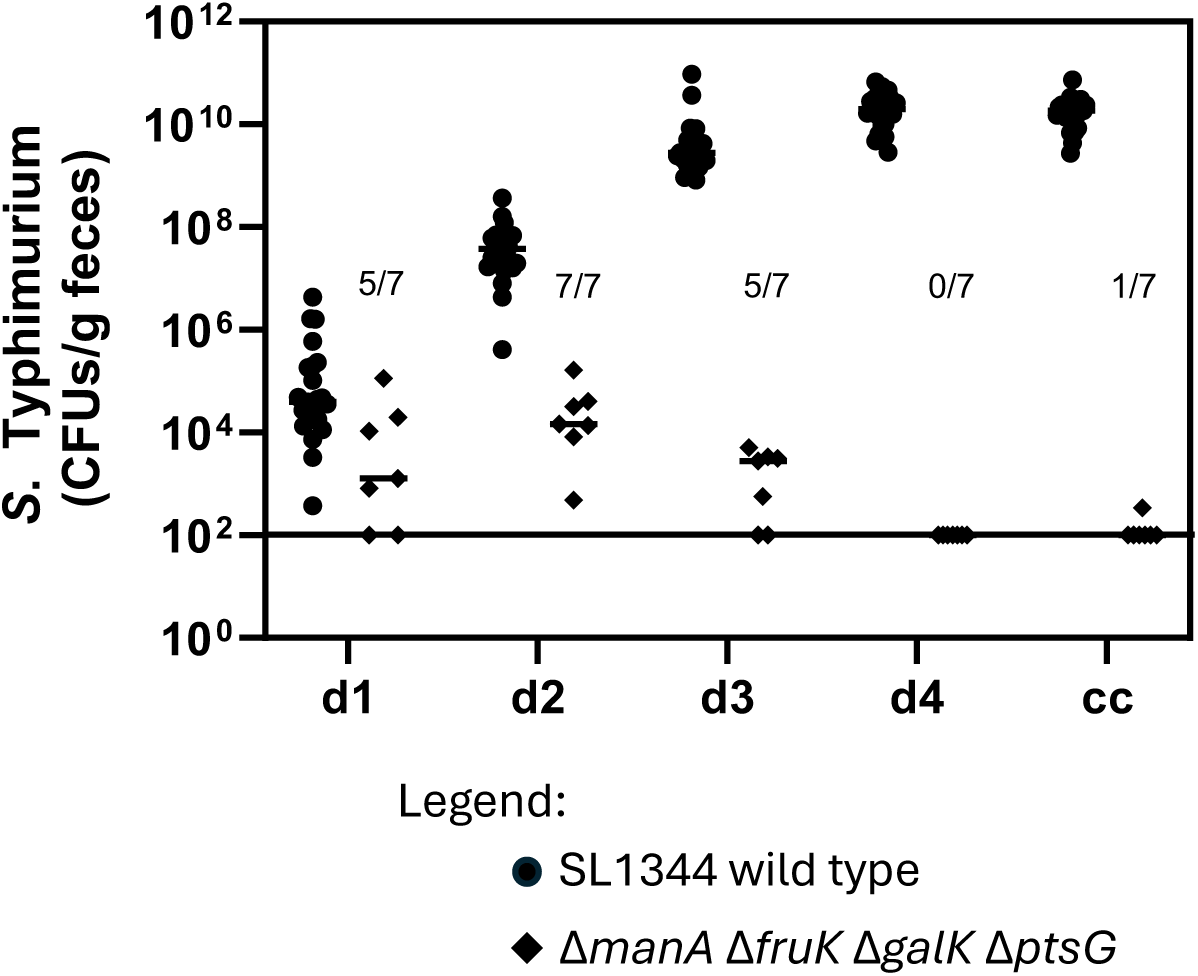
The Δ*manA* Δ*fruK* Δ*galK* Δ*ptsG* mutant is displaced by the wild type in OligoMM^12^. Competitive infection of C57BL/6J mice associated with the OligoMM^12^ microbiota with SL1344 wild type (circle; mice n=23) and Δ*manA* Δ*fruK* Δ*galK* Δ*ptsG* quadruple mutant (diamond; mice n=7, at least two independent experiments). The bacterial loads are plotted in CFU per gram of feces. The detection limit is indicated, and the numbers above show how many samples were below the detection limit. SL1344 wild type actively displaced the quadruple carbohydrate mutant in a mouse model with an intact microbiota.

**Figure S5:**
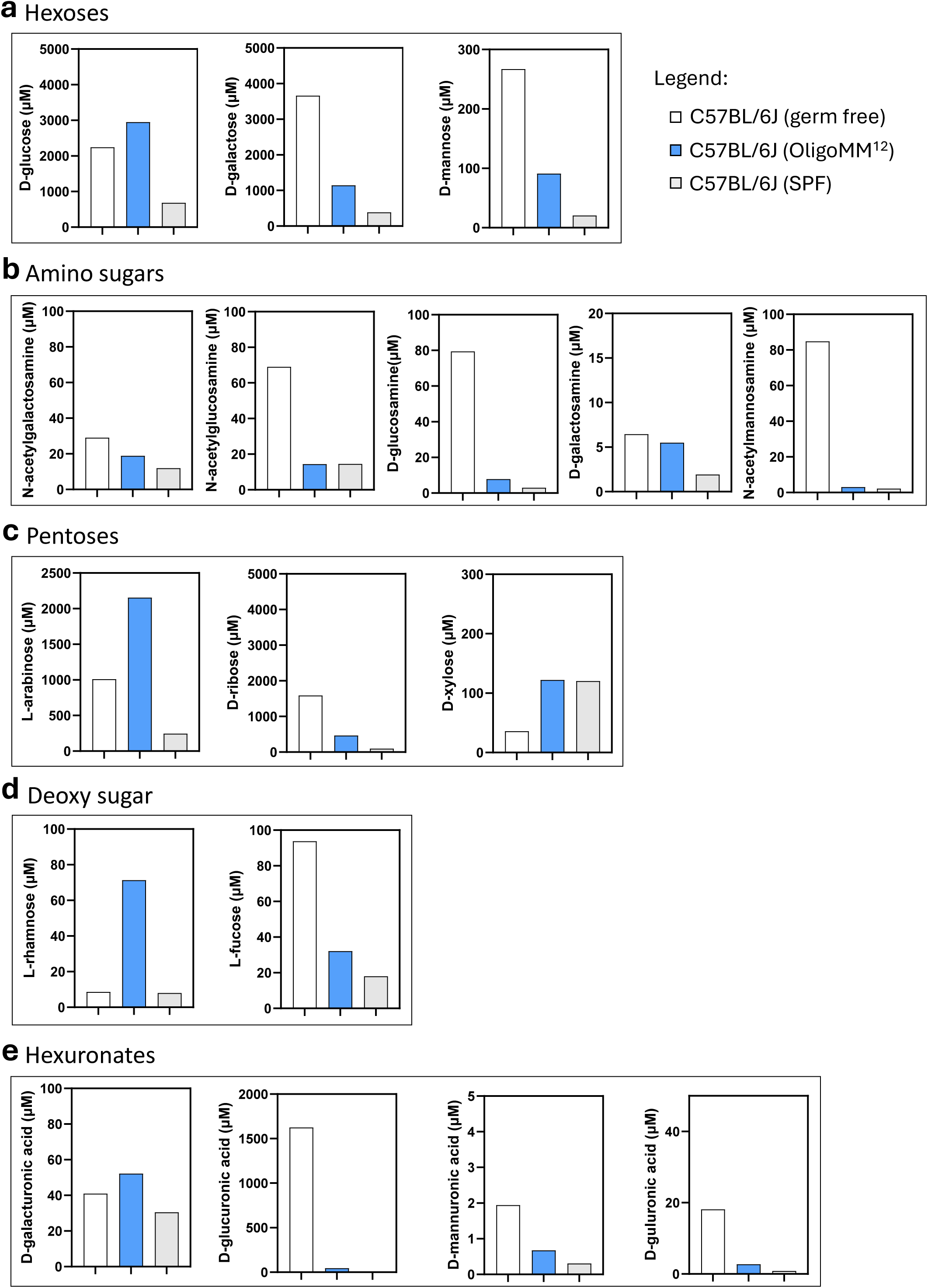
Monosaccharide quantification by LC-MS. The monosaccharides are presented in separate blots, showing the median amount on a linear y-axis scale. The data of C57BL/6J germ-free (mice n=5) and OligoMM^12^ models (mice n=5) were compared to the previously published dataset of SPF C57BL/6J (mice n=5) (Nguyen et al, 2024). **A** Hexoses, including D-glucose, D-galactose, and D-mannose. **B** Amino sugars, consisting of N-acetylgalactosamine, N-acetylglucosamine, D-glucosamine, D-galactosamine, and N-acetylmannosamine. **C** Pentoses, including L-arabinose, D-ribose, and D-xylose. **D** Deoxy sugars, consisting of L-rhamnose, and L-fucose. **E** Hexuronates, including D-galacturonic acid, D-glucuronic acid, D-mannuronic acid, and D-guluronic acid.

**Figure S6:**
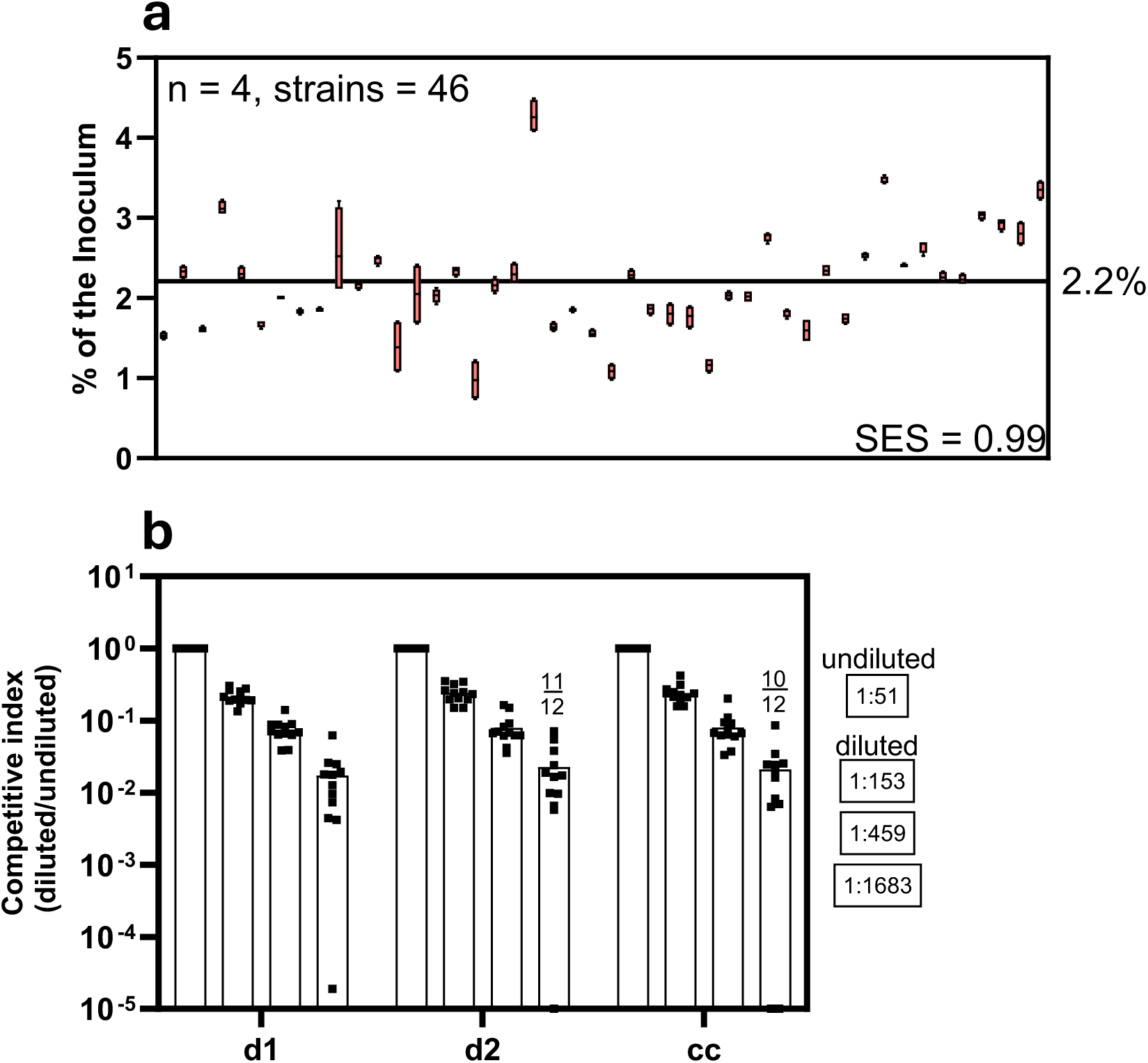
Strain distribution of the infection mixture and detection limit. **A** The Shannon evenness score (SES) was calculated for all 46 SL1344 strains within the pool, excluding the wild-type titration standards. The y-axis indicates the proportion of each SL1344 mutant within the inoculum across four distinct inocula used in the mouse experiments. The average abundance across all mutants in the inoculum should be 2.2%. The SL1344 mutant pool achieves a SES of 0.99. **B** The competitive index was plotted for the titration standards normalized to an undiluted SL1344 wild type. The dilution ratios are indicated on the right side of the diagram. The number above the bar shows how many samples are within detection limit. We still have excellent resolution at 10^-3^; however, this can vary due to fluctuations in sequencing depth.

